# Strategies for Partitioning Clock Models in Phylogenomic Dating: Application to the Angiosperm Evolutionary Timescale

**DOI:** 10.1101/144287

**Authors:** Charles S. P. Foster, Simon Y. W. Ho

## Abstract

Evolutionary timescales can be inferred from molecular sequence data using a Bayesian phylogenetic approach. In these methods, the molecular clock is often calibrated using fossil data. The uncertainty in these fossil calibrations is important because it determines the limiting posterior distribution for divergence-time estimates as the sequence length tends to infinity. Here we investigate how the accuracy and precision of Bayesian divergence-time estimates improve with the increased clock-partitioning of genome-scale data into clock-subsets. We focus on a data set comprising plastome-scale sequences of 52 angiosperm taxa. There was little difference among the Bayesian date estimates whether we chose clock-subsets based on patterns of among-lineage rate heterogeneity or relative rates across genes, or by random assignment. Increasing the degree of clock-partitioning usually led to an improvement in the precision of divergence-time estimates, but this increase was asymptotic to a limit presumably imposed by fossil calibrations. Our clock-partitioning approaches yielded highly precise age estimates for several key nodes in the angiosperm phylogeny. For example, when partitioning the data into 20 clock-subsets based on patterns of among-lineage rate heterogeneity, we inferred crown angiosperms to have arisen 198–178 Ma. This demonstrates that judicious clock-partitioning can improve the precision of molecular dating based on phylogenomic data, but the meaning of this increased precision should be considered critically.

## Introduction

Evolutionary timescales can be estimated from molecular sequence data using phylogenetic methods based on the molecular clock. In practice, most data sets exhibit substantial rate heterogeneity among lineages. These ‘lineage effects’ can be caused by variation in life-history traits, generation time, or exposure to mutagens (Smith and Donoghue 2008; Gaut et al. 2011; Lanfear et al. 2013). Among-lineage rate variation can be taken into account using Bayesian relaxed-clock models, in which the rates can be assumed to be either correlated between neighbouring branches (Thorne et al. 1998; Kishino et al. 2001) or drawn independently from a chosen distribution (Drummond et al. 2006; Rannala and Yang 2007).

A number of factors can cause rates to vary across loci in the genome (Wolfe et al. 1987). These ‘gene effects’ can be taken into account by allowing each locus to have a distinct relative rate. Less certain is the best way to deal with interactions between gene effects and lineage effects, which can be caused by differences in selective pressure and other processes (Gaut et al. 2011). In this case, the extent and patterns of among-lineage rate heterogeneity vary across genes or other subsets of the data. This form of rate variation can be captured by assigning separate clock models to different subsets of the data (Ho and Duchêne 2014), a process that we refer to here as clock-partitioning.

Appropriate clock-partitioning can improve the precision of Bayesian date estimates (as measured by the associated 95% credibility intervals), but it is rarely done in practice. This is also despite widespread adoption of partitioning schemes for substitution models (Lanfear et al. 2012). The most likely explanation is that the use of clock-partitioning in Bayesian phylogenetics greatly increases the risk of overparameterization, and thus to reduced Markov chain Monte Carlo performance. Overparameterization has been previously addressed in light of the bias-variance trade-off, which is well established in statistical theory (Burnham and Anderson 2003). Compared with a complex, parameter-rich model, a simple model that underfits data is expected to have low accuracy (high bias) but high precision (low variance). Conversely, a parameter-rich model that overfits the data is likely to have higher accuracy, but this comes at the cost of reduced precision. The best model is an intermediate one that simultaneously maximizes accuracy and precision (Wertheim et al. 2010)

It is useful to consider the bias-variance trade-off in the context of molecular dating with partitioned clock models. Patterns of among-lineage rate variation are likely to differ across genes (Muse and Gaut 1994), so increasing the number of relaxed clocks will better capture these patterns of rate heterogeneity and should lead to more accurate age estimates (Duchêne and Ho 2014). However, each clock-subset has parameters that need to be estimated, including a distinct set of branch rates. As a consequence, increasing the degree of clock-partitioning should lead to a widening of the posterior distributions of parameters.

Contrary to the expectations of the bias-variance trade-off, increasing the degree of clock-partitioning tends improve the precision of Bayesian age estimates (Zhu et al. 2015). One possible explanation for this lies in the treatment of the uncertainty in the estimates of genetic branch lengths. The accuracy and precision of evolutionary rate estimates depend on the accurate inference of branch lengths (in substitutions per site). In the case of molecular dating, branch rates for each clock-subset are combined with node times to give the branch lengths. Therefore, as the number of clock-subsets increases, the node times in the chronogram are estimated from an increasing number of data points, leading to increasing precision. Although branch-length estimation generally improves as the amount of sequence data increases, branch lengths can be estimated with reasonable accuracy even with fairly small amounts of sequence data (Yang and Rannala 2006). This suggests that for a data set of a (large) fixed size, increasing the number of clock-subsets should lead to improved precision in divergence-time estimates until the amount of sequence data in each clock-subset decreases to a critical point.

Zhu et al. (2015) explain this phenomenon in their ‘finite sites’ theory, although they use the term ‘loci’ to refer to clock-subsets. Even with sequences of infinite length, there will still be uncertainty in the age estimates, corresponding to the uncertainty in the fossil calibrations (“infinite data limit”; Yang and Rannala 2006; dos Reis and Yang 2013). As the number of clock-subsets (*L*) increases, the finite-sites theory suggests that the uncertainty in age estimates decreases to the infinite-data limit at the rate of 1/*L* (Zhu et al. 2015). This property has important consequences for analyses of genome-scale data sets, whereby many genes are analysed concurrently. Therefore, it is important that both the finite-sites theory and the bias-variance trade-off are tested comprehensively on a genome-scale data set with clock-partitioning.

Persistent uncertainty in molecular date estimates is perhaps best exemplified by studies of the origins of flowering plants (angiosperms) (Foster 2016). The earliest unequivocal angiosperm fossils are tricolpate pollen grains from the Barremian–Aptian boundary, from approximately 125.9 million years ago (Ma) (Hughes 1994). Older pollen grains from the Hauterivian provide some evidence of crown-group angiosperms, and are usually accepted as belonging to this group, albeit with less confidence than for the tricolpate pollen grains (Herendeen et al. 2017). Patterns of diversification in the broader fossil record suggest that angiosperms are unlikely to have arisen much earlier than this time (Magallón et al. 2015). The majority of molecular dating analyses tell a vastly different story, with most recent analyses inferring an origin within the Triassic (Foster et al. 2017). Additionally, the uncertainty surrounding the age of the angiosperm crown node is large, often spanning an interval of many tens of millions of years, unless strong age constraints are placed on the node. Improving the accuracy and precision of estimates of the age of crown angiosperms thus represents a key goal of molecular dating.

In this study, we use a Bayesian phylogenetic approach to investigate the impact of clock-partitioning on the precision of divergence-time estimates. We also investigate whether the criteria used to assign genes to different clocks has an impact on estimation error. To do so, we infer the evolutionary timescale of angiosperms using a plastome-level data set. In analyses with clock-partitioning schemes comprising up to 20 clock-subsets, we allocate genes to clock-subsets based on patterns of among-lineage rate heterogeneity or relative substitution rate, or through random assignment. In all cases, we confirm that increasing the degree of clock-partitioning can lead to vast improvements in the precision of Bayesian date estimates.

## Materials and Methods

### Data Sets and Clock-Partitioning

We obtained full chloroplast genome sequences for 52 angiosperm taxa and two gymnosperm outgroup taxa from GenBank (supplementary table S1, Supplementary Material online). Each angiosperm taxon was chosen to represent a different order, with our sampling designed to include as many as possible of the 63 angiosperm orders recognized by the Angiosperm Phylogeny Group (2016). We extracted all 79 protein-coding genes from the chloroplast genomes, although some genes were missing from some taxa. We initially translated all genes into amino acid sequences using VirtualRibosome (Wernersson 2006) and aligned them using MAFFT v7.305b (Katoh and Standley 2013). We then translated the aligned amino acid sequences back into nucleotide sequence alignments using PAL2NAL (Suyama et al. 2006), made manual adjustments, and filtered out any sites in the alignment at which a gap was present in ≥80% of the taxa. Our total core data set consisted of 68,790 nucleotides, of which only 7.54% sites were gaps or missing data (see supplementary file S1, Supplementary Material online).

Our primary strategy for clock-partitioning based on patterns of among-lineage rate heterogeneity was to analyse the genes using ClockstaR v2 (Duchêne et al. 2014). ClockstaR takes predefined subsets of the data, along with the estimated gene tree for each subset, and determines the optimal clock-partitioning scheme for the data set. This involves identifying the optimal number of clock-subsets (*k*), as well as the optimal assignment of the data subsets to each of these clock-subsets. Comparison of clock-partitioning schemes is done by comparing the patterns of among-lineage rate heterogeneity across the gene trees and clustering the gene trees according to the gap statistic (Gap_*k*_) (Tibshirani et al. 2001). Additionally, ClockstaR can determine the optimal clock-partitioning scheme for any value of *k*. In our case, each of the 79 protein-coding genes was considered as a separate data subset for the ClockstaR analysis.

ClockstaR requires all data subsets to share the same tree topology. Since the chloroplast genome does not typically undergo recombination (Birky 1995), all of its genes should share the same topology. Therefore, we first inferred the phylogeny for the concatenated data set using maximum-likelihood analysis in IQ-TREE v1.50a (Nguyen et al. 2015), with node support estimated using 1000 bootstrap replicates with the ultrafast bootstrapping algorithm (Minh et al. 2013). We partitioned the data set by codon position using the edge-linked partition model (Chernomor et al. 2016), and implemented the GTR+Γ_4_ model of nucleotide substitution for each subset. The best-scoring tree was very similar to previous estimates of the angiosperm phylogeny based on chloroplast data (Moore et al. 2010; Soltis et al. 2011), and we found strong support for most nodes in the tree (supplementary fig. S1, Supplementary Material online). We used this tree for ClockstaR and optimized the branch lengths for each gene alignment. Finally, we determined the optimal value of *k*, and then created 12 clock-partitioning schemes using the optimal assignment of genes to clock-subsets for values of *k* from 1 to 10, 15, and 20 (“*P_CSTAR_*” schemes). We use the partitioning along medoids (PAM) algorithm, described by Kaufman and Rousseeuw (2009).

As a means of comparison with the ClockstaR partitioning schemes, we also chose clock-partitioning schemes based on relative substitution rates across genes (dos Reis et al. 2012). To do so, we focused on a subset of 20 taxa for which sequences of all 79 protein-coding genes were available (supplementary table S1, Supplementary Material online). We then analysed each gene using maximum likelihood in IQ-TREE, in each case partitioning by codon position and implementing the GTR+Γ_4_ model of nucleotide substitution for each codon position. Using the tree lengths as a proxy for the overall substitution rate of each gene, we created 11 partitioning schemes based on relative rates of substitution (“*P_RATE_*” schemes), in which we assigned genes to clock-subsets for values of *k* from 2 to 10, 15, and 20.

For an additional form of comparison, we generated clock-partitioning schemes with genes randomly allocated to clock-subsets. Genes were randomly sampled without replacement in R v3.3.2 (R Core Team 2016) and assigned to clock-subsets for values of *k* from 2 to 10, 15, and 20. We repeated this process three times, resulting in a total of 33 clock-partitioning schemes in which genes were randomly assigned to clock-subsets (“*P_RAND_*” schemes).

### Molecular Dating

We inferred the evolutionary timescale using MCMCTREE in PAML v4.8 (Yang 2007) with the GTR+Γ_4_ model of nucleotide substitution. A key requirement of MCMCTREE is a fixed tree topology, so we used the best-scoring tree that we estimated from the total concatenated data set using IQTREE. We primarily analysed our data sets with the UCLN relaxed clock (Drummond et al. 2006; Rannala and Yang 2007), but replicated all analyses to check for any differences under the ACLN relaxed clock (Thorne et al. 1998; Kishino et al. 2001).

We estimated the overall substitution rate for each clock-partitioning scheme by running baseml under a strict clock, with a single point calibration at the root. We then used this estimate to select the shape (*α*) and scale (*β*) parameters for the gamma-Dirichlet prior on the overall substitution rate across loci in the MCMCTREE analysis according to the formulae *α* = (*m/s*)^2^ and *β* = *m/s*^2^, where *m* and *s* are the mean and standard deviation of the substitution rate, respectively. For all analyses, we set the shape and scale parameters for the gamma-Dirichlet prior on rate variation across branches to 1 and 3.3, respectively. The posterior distribution of node ages was estimated with Markov chain Monto Carlo sampling, with samples drawn every 10^3^ steps across a total of 10^7^ steps, after a discarded burn-in of 10^6^ steps. We ran all analyses in duplicate to assess convergence, and confirmed sufficient sampling by checking that the effective sample sizes of all parameters were above 200.

We repeated the MCMCTREE analysis for all *P_CSTAR_, P_RATE_*, and *P_RAND_* schemes. An advantage of MCMCTREE is the option to use approximate likelihood calculation, which is much faster than full likelihood calculation (Thorne et al. 1998; dos Reis and Yang 2011). However, this precludes the calculation of marginal likelihoods using path sampling and similar methods, which require the full likelihood to be computed. Instead, we compared the means and 95% credibility intervals of the posterior estimates of divergence times across our partitioning strategies. We chose to focus on six nodes in the angiosperm phylogeny: the crown groups of all angiosperms, magnoliids, monocots, eudicots, campanulids, and Liliales. The first four of these were chosen because they define major clades in the angiosperm phylogeny. The other two nodes were chosen because they do not have explicit fossil-based calibration priors.

### Fossil Calibrations

Calibrations are the most important component of Bayesian molecular dating, with critical impacts on posterior estimates of divergence times. Therefore, we selected a set of 23 calibration priors primarily based on recent studies that carefully considered the phylogenetic affinities of angiosperm fossils (table 1). We also applied two calibration priors to the gymnosperm outgroup. Fossils can strictly only provide a minimum age for the divergence of lineages from their common ancestor, so we chose to implement fossil calibrations primarily as uniform distributions with soft bounds. This approach assigns an equal prior probability for all ages between specified minimum and maximum ages, with a 2.5% probability that the age surpasses each bound (Yang and Rannala 2006).

**Table 1.**
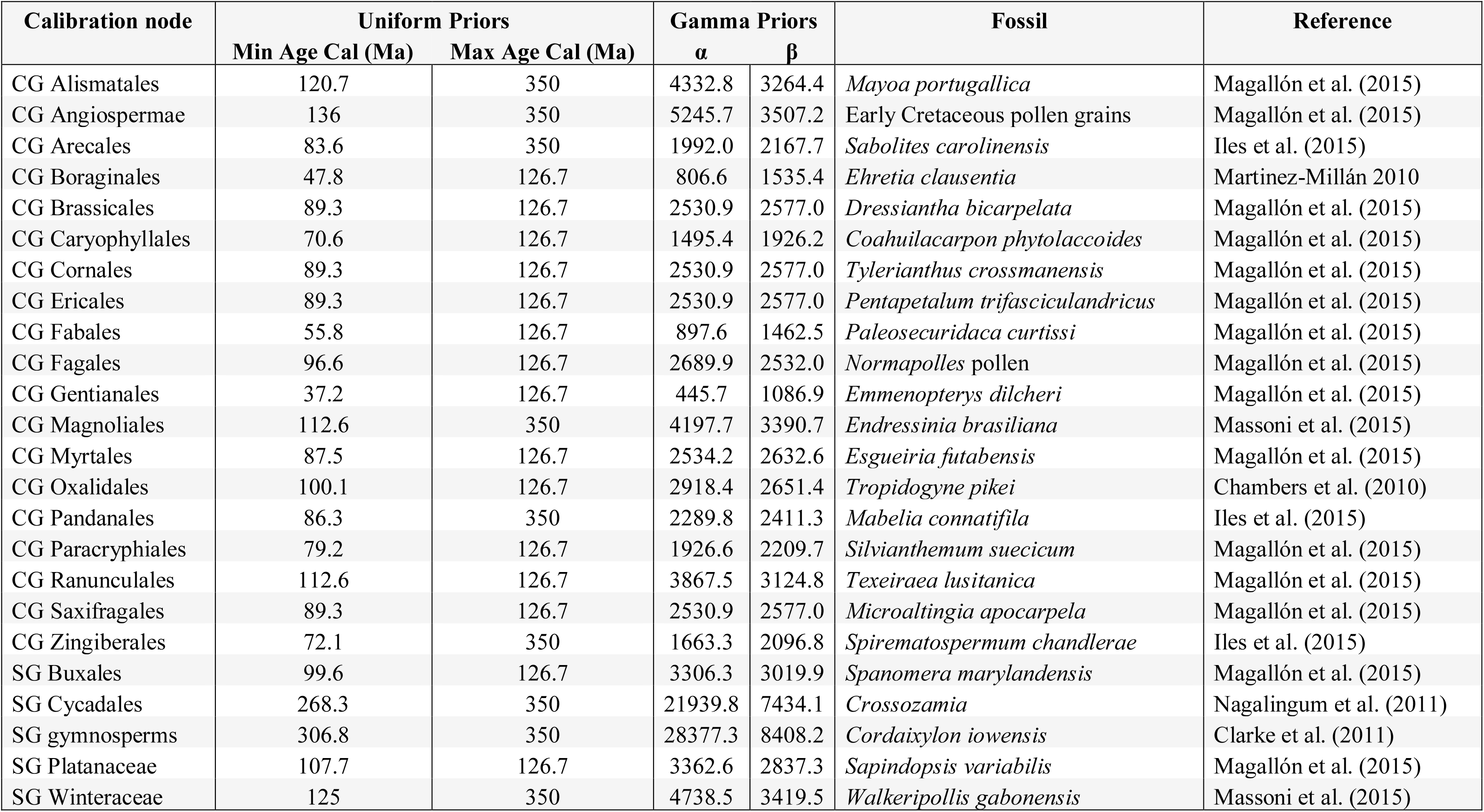
—The calibration priors used within this study to estimate the angiosperm evolutionary timescale. “CG” and “SG” refer to the crown and stem groups, respectively, of the clade of interest.

We implemented two maximum age constraints: (i) 350 Ma for the divergence between angiosperms and gymnosperms (the root), a well accepted upper bound for this divergence (Foster et al. 2017); and (ii) 126.7 Ma for the origin of crown eudicots, corresponding to the upper bound of the Barremian–Aptian boundary (reviewed by Massoni et al. 2015a). The latter constraint is widely used and is justified by the complete absence of tricolpate pollen before the latest Barremian, yet some molecular dating results have suggested an earlier origin for eudicots (Smith et al. 2010; Foster et al. 2017; Zeng et al. 2017). Ranunculales, one of the earliest-diverging eudicot orders, has a fossil record dating back to the late Aptian/early Albian. Therefore, implementing the eudicot maximum constraint results in a strong prior being placed on crown-group eudicots appearing between ~126.7–112.6 Ma. As a result, including the eudicot maximum constraint leads to the eudicot crown node being a useful example of a heavily constrained node for downstream comparisons of the uncertainty in posterior age estimates.

For comparison, we also performed analyses with our *P_CSTAR_* schemes using gamma calibration priors and the UCLN relaxed clock. In this case, the mean of each gamma prior was set to the age of each fossil +10%, with an arbitrary standard deviation of 2 (Table 1). This effectively brackets the age estimates of calibrated nodes within a very narrow interval. In such a calibration scheme, the precision of age estimates is not expected to improve substantially with increased clock-partitioning.

## Results

### Angiosperm Evolutionary Timescale

Our ClockstaR analysis identified the optimal value of *k* to be 1, suggesting that a single pattern of among-lineage rate heterogeneity is shared across protein-coding genes from the chloroplast genomes. However, despite *k*=1 being optimal, the values of the gap statistic were still higher for all values of *k*>5 (figure 1). Based on our analysis using the optimal clock-partitioning scheme (*k*=1) and the UCLN relaxed clock, we estimated the time to the most recent common ancestor of angiosperms to be 196 Ma (95% credibility interval 237–161 Ma; supplementary fig. S2, Supplementary Material online). We inferred that crown magnoliids first appeared 171– 115 Ma, and that crown monocots arose contemporaneously, 167–120 Ma. Crown eudicots were inferred to have arisen 128–124 Ma, with this precise estimate reflecting the strong calibration prior placed upon this node. Finally, our estimates for the time to the most recent common ancestors of campanulids and Liliales were 101–91 Ma and 108–91 Ma, respectively.

**Fig. 1.**
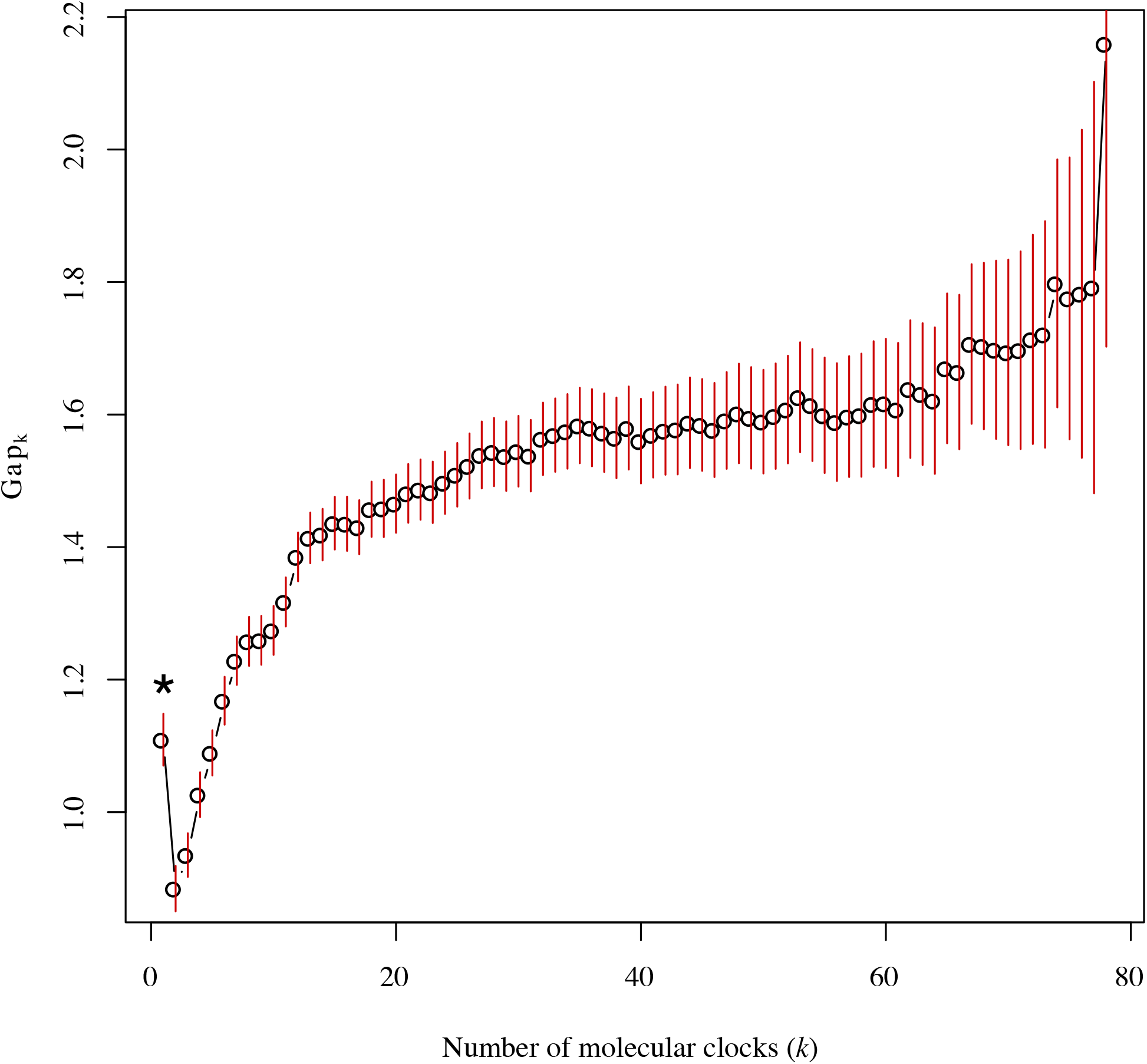
Gap statistic values for different numbers of clock-subsets (*k*) for the plastome-scale angiosperm data set, inferred using partitioning along medoids in ClockstaR. The asterisk indicates the optimal number of clock-subsets.

The true age of crown angiosperms is unknown, so we cannot assess the absolute accuracy of our date estimates. Instead, we consider the consistency of mean age estimates across analyses (Hillis 1995). The mean age estimates for all crown angiosperms, magnoliids, and monocots varied slightly across values of *k* from 1 to 3, but estimates remained stable across all other values of *k*. Mean age estimates for crown eudicots only varied by approximately 2 myr across all values of *k*. Mean age estimates for crown Liliales were stable across all clock-partitioning schemes. However, mean estimates for crown campanulids steadily declined by approximately 10–15 myr as the number of loci increased. We observed the same broad trends in accuracy for all nodes of interest when using the ACLN relaxed clock, although mean age estimates were consistently slightly younger than in analyses with the UCLN relaxed clock. In our analyses with the *P_CSTAR_* schemes and with gamma calibration priors, mean age estimates for crown angiosperms steadily increased with increasing numbers of clock-subsets, but the mean estimates were stable for all other nodes of interest.

### Precision in Estimates of Divergence Times

We focus first on our results when using the UCLN relaxed clock, uniform calibration priors, and with clock-partitioning according to ClockstaR. We report improvements in the precision of node-age estimates by calculating the decrease in 95% CI width, which we standardized by dividing by the posterior mean. The optimal clock-partitioning scheme was inferred to be *k*=1, matching the results of previous analyses (Duchêne et al. 2016). However, increasing the number of clock-subsets generally led to large increases in the precision of node-age estimates. The impact of this is perhaps most striking in the inferred age of crown angiosperms. Increasing the number of clock-subsets from *k*=1 to *k*=2 led to a reduction in statistical fit (figure 1), but also reduced the width of the 95% CI for the inferred age of crown angiosperms from 77 myr to 46 myr (an improvement in precision of 35.4%). Greater clock-partitioning led to further improvement in precision (figure 2). For example, implementing a clock-partitioning scheme with *k*=20 reduced the width of the 95% CI for the inferred age of crown angiosperms to only 20 myr, representing a 73.1% improvement in precision. However, the rate of improvement in precision declined rapidly for increasing numbers of clock-subsets (figure 2).

**Fig. 2.**
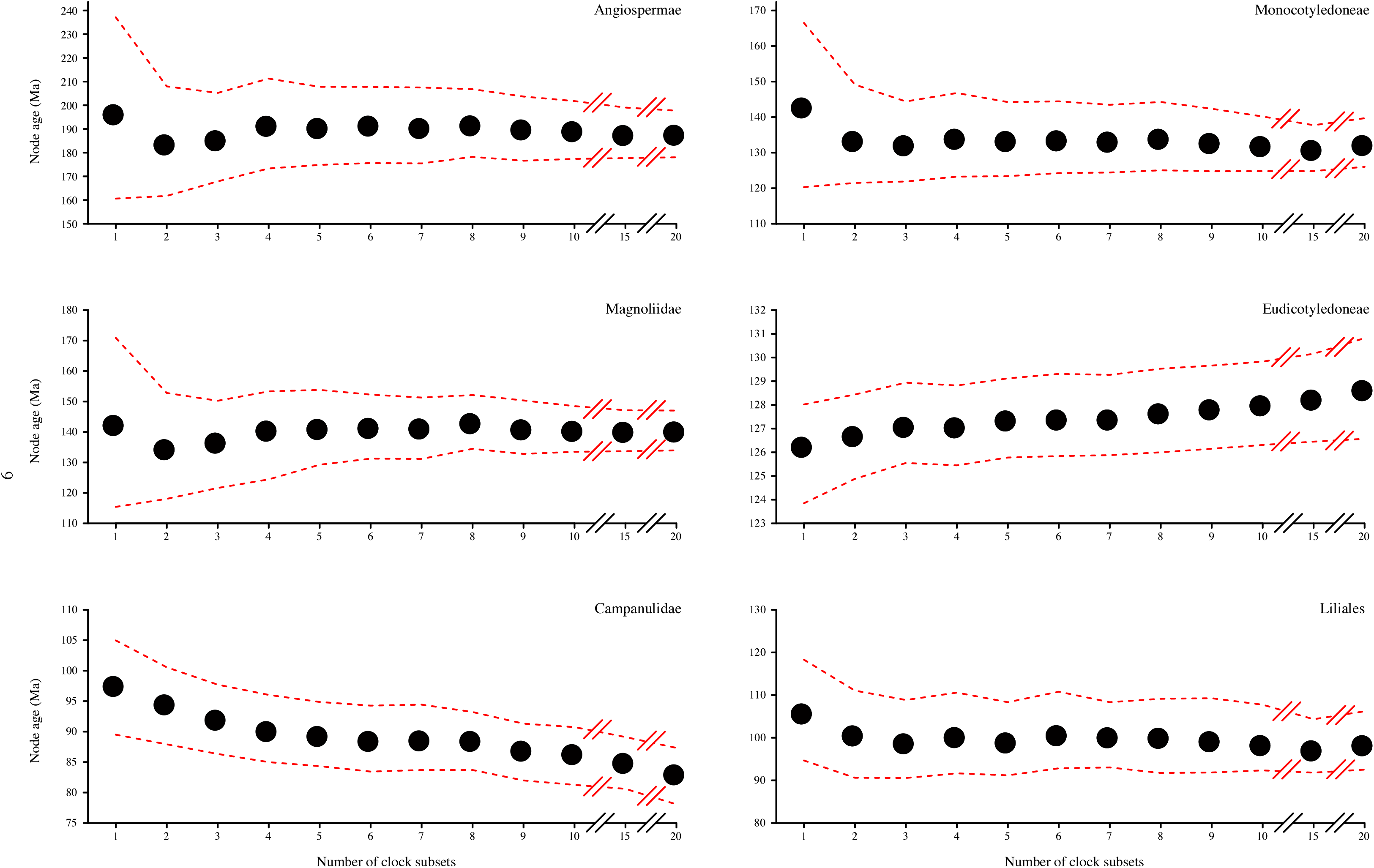
Mean age inferences and associated 95% credibility intervals for six nodes in the angiosperm phylogeny with increasing numbers of clock-subsets (*k*), as inferred using an autocorrelated lognoπnal relaxed clock, clock-partitioning according to the optimal schemes identified in ClockstaR, and uniform calibration priors.

An improvement in precision with the number of clock-subsets can also be observed in the age estimates for both magnoliids and monocots. For example, increasing *k* from 1 to 20 results in respective increases of 76.1% and 68% in precision in the age estimates for crown magnoliids and crown monocots (figure 2). When considering the nodes corresponding to the crown groups of campanulids and Liliales, a similar trend can be observed, albeit with a less drastic increase in precision. Increasing the number of clock-subsets led to 29.7% and 37.7% increases in precision for the crown groups of campanulids and Liliales, respectively. However, there is a vastly different trend in the age estimate for crown eudicots. In this case, the age estimate for *k*=1 is already precise (95% credibility interval: 128–124 Ma) and increasing the number of clock-subsets actually led to a slight decrease in precision of 0.02%.

Compared with the *P_CSTAR_* clock-partitioning schemes, very similar trends in precision were observed for both the *P_RATE_* scheme (figure 3) and *P_RAND_* scheme (figure 4). The only differences were that there was less variation in mean age estimates for smaller values of *k* compared with the ClockstaR partitioning scheme, and standardized improvements in precision were consistently slightly greater (supplementary table S2, Supplementary Material online). For example, the widths of the 95% CIs, and the mean age estimates, declined monotonically in both classes of clock-partitioning schemes.

**Fig. 3.**
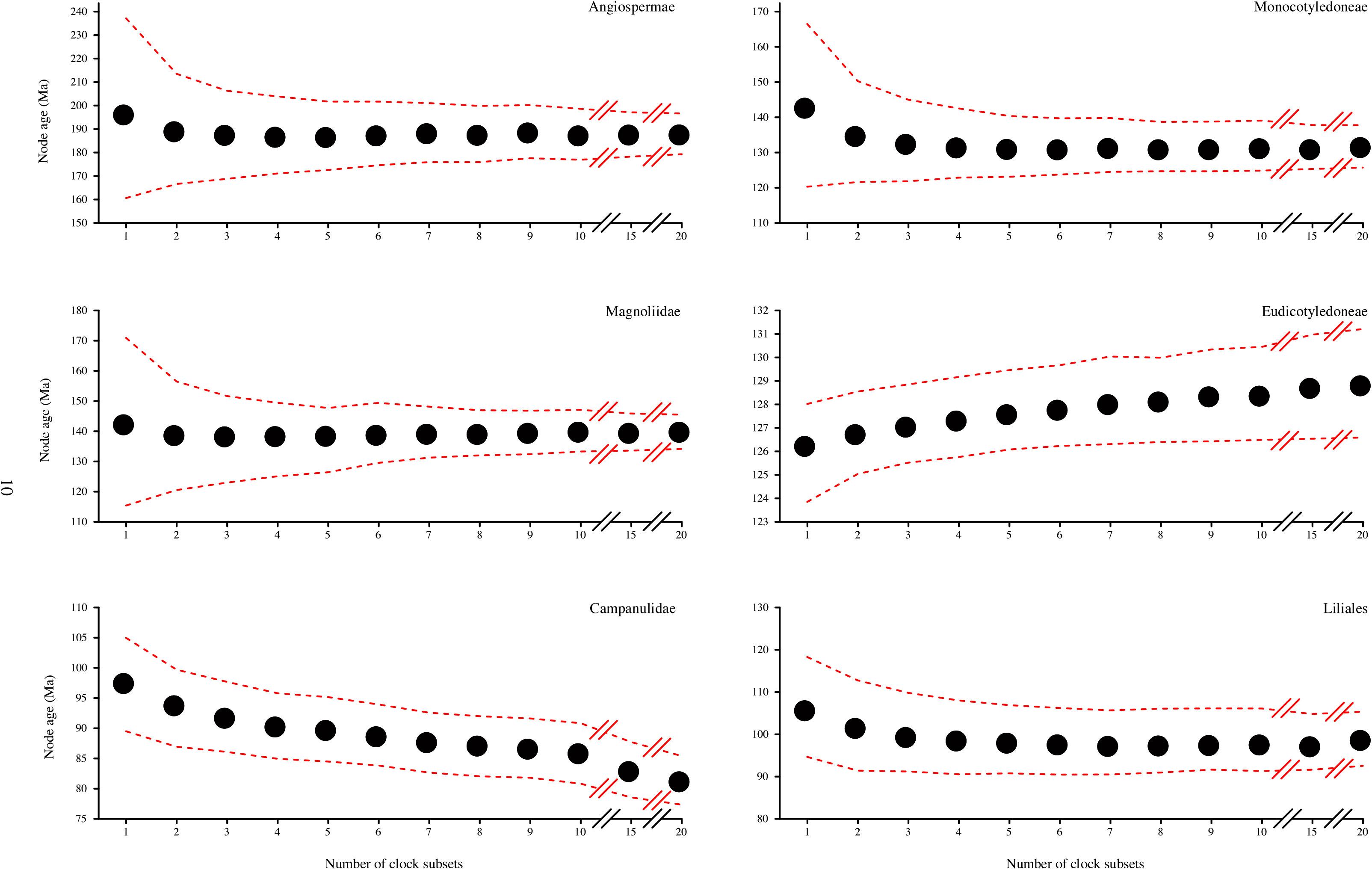
Mean posterior age estimates and associated 95% credibility intervals for six nodes in the angiosperm phylogeny with increasing numbers of clock-subsets (*k*), as inferred using an uncorrelated lognoπnal relaxed clock, clock-partitioning according to relative rates of substitution, and uniform calibration priors.

**Fig. 4.**
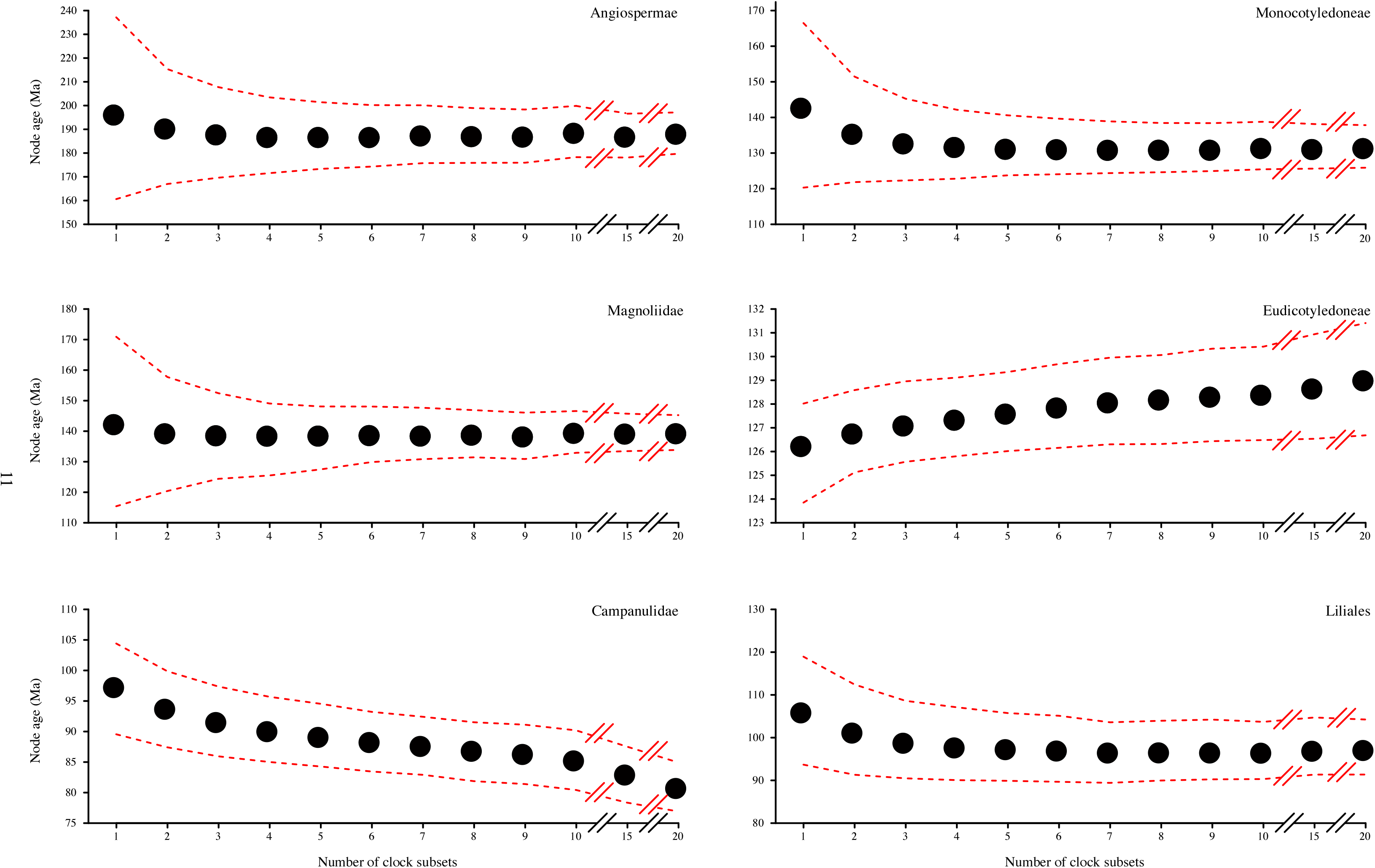
Mean posterior age estimates and associated 95% credibility intervals for six nodes in the angiosperm phylogeny with increasing numbers of clock-subsets (*k*), as inferred using an uncorrelated lognormal relaxed clock, clock-partitioning according to random assignment of genes to clock subsets, and uniform calibration priors. The estimates presented here are the averages of three random assignments of genes to clock-subsets for each value of *k*.

We observed the same broad trends across all clock-partitioning schemes when using the ACLN relaxed clock. With increasing numbers of clock-subsets, the uncertainty in age estimates rapidly decreased, with the exception of the age estimate for the eudicot crown node. Even with *k*=1, however, the precision of the age estimates was much greater than in the corresponding analysis with the UCLN relaxed clock. For example, when implementing the *P_CSTAR_* clock-partitioning schemes, the 95% credibility interval of the age estimate for crown angiosperms spanned 77 myr when using the UCLN relaxed clock, but only 59 myr when using the ACLN relaxed clock. Additionally, age estimates for crown eudicots became less precise as the degree of clock-partitioning increased. We observed the same trend for the other nodes of interest across analyses, and the apparent limit to uncertainty appeared to be reached much more rapidly than with the UCLN relaxed clock (supplementary fig. S3–S5, supplementary table S2, Supplementary Material online).

When using highly informative gamma calibration priors in our additional analyses of the *P_CSTAR_* schemes, we found that for the crown groups of angiosperms, monocots, and magnoliids, the increases in precision with greater clock-partitioning were much lower than with uniform calibration priors (supplementary fig. S6 and supplementary table S2, Supplementary Material online). For example, an improvement of only 18.5% occurred in the precision of the age estimate for crown angiosperms. The opposite trend occurred for the crown nodes of eudicots, campanulids and Liliales. When implementing uniform calibration priors, greater clock-partitioning led to either no change or decreases in precision for age estimates of crown-group eudicots, but when using gamma calibration priors the precision improved by 36% with greater clock-partitioning. For crown-group Liliales, increasing *k* from 1 to 20 led to a 64.3% increase in the precision of age estimates, the greatest improvement of all six key nodes. However, it is worth noting that our age estimates for all six nodes of interest were very precise even when *k*= 1. Therefore, in terms of absolute time units, there was generally little improvement in precision with increasing numbers of clock-subsets.

## Discussion

The primary aim of the present study was not to provide a novel estimate for the angiosperm evolutionary timescale, but it is still useful to consider our results in the context of previous estimates. Our inferred origin for crown-group angiosperms in the late Triassic to early Jurassic is consistent with most modern molecular dating estimates (Bell et al. 2010; Magallón 2010; Clarke et al. 2011; Zeng et al. 2014; Beaulieu et al. 2015; Foster et al. 2017). Similarly, our age estimate for crown magnoliids of 171–115 Ma is very similar to a previous estimate of 179–127 Ma based on the most comprehensive molecular dating analyses of Magnoliidae (Massoni et al. 2015a). Our estimate of 167–120 Ma for the age of crown monocots is compelling, because a recent study of monocots using the fossilized-birth-death model inferred a very similar age of 174–134 Ma (Eguchi and Tamura 2016). Our age estimate for crown eudicots of 128–124 Ma suggests that there was not enough signal within the data to overcome the strong calibration priors placed upon this node. Finally, although our age estimate for the appearance of crown campanulids 101–91 Ma is very similar to those of recent studies (Magallón et al. 2015; Foster et al. 2017), our age estimate of 108–91 Ma for the time to the most recent common ancestor of Liliales was slightly younger than recent estimates.

The goal of all molecular dating studies is to estimate the evolutionary timescale with a useful degree of precision and accuracy. We demonstrated that increasing the degree of clock-partitioning leads to increasingly precise age estimates, as predicted by the finite-sites theory (Zhu et al. 2015). Additionally, clock-partitioning schemes based on patterns of among-lineage rate heterogeneity or relative substitution rates did not have any measurable advantage over randomly assigning genes to clock-subsets, at least in terms of the accuracy and precision of the resulting estimates of divergence times. The near-identical patterns of precision across all clock-partitioning schemes stands in contrast with previous suggestions that the assignment of genes to clock-subsets is more important than the number of clock-subsets (Duchêne and Ho 2014).

Our results demonstrate that to improve the precision of age estimates, one could simply increase the degree of clock-partitioning by assigning genes to an arbitrarily large number of clock-subsets, until the marginal benefit of increasing the number of clocks is close to zero (Zhu et al. 2015). An obvious consequence of this is that one must consider whether such an increase is desirable or biologically meaningful. If there is evidence that a data set conforms to a single pattern of rate variation among lineages, an increase in precision from clock-partitioning is not justifiable because the clock-subsets do not constitute independent realizations of the process of rate variation (Zhu et al. 2015). Our analysis using ClockstaR indicates that within our data set, all genes exhibit the same pattern of rate heterogeneity among lineages, such that they should be analysed using a single clock model. In this case, increasing the degree of clock-partitioning leads to a model that overfits the data, does not appear to accurately predict the data, and is insensitive to the sampled data. Normally this would be expected to occur when a model underfits the data, but the increasing sets of “independent” branch-rate estimates for each clock-subset ensure that estimates of node times remain precise.

The uncertainty in posterior divergence times can be divided into three components: (i) uncertainty in branch lengths due to limited sequence length (*N*); (ii) among-lineage rate variation for each clock-subset, as well as the evolutionary rate variation among clock-subsets; and (iii) uncertainty in fossil calibrations (Zhu et al. 2015). If *L* is large, then the uncertainty caused by limited sequence length approaches zero at the rate of 1/*N*. Additionally, the uncertainty attributable to the second component approaches zero at the rate of 1/*L*. As *N*→∞ and *L*→∞, the uncertainty in divergence-time estimates should be wholly attributable to uncertainty in the fossil calibrations (Zhu et al. 2015). For a data set of fixed size, such as our angiosperm data set, increasing *L* will reduce *N*, and vice versa. We found that partitioning the data set into increasing numbers of clock-subsets led to improvements in precision, which implies that increasing *L* has a larger impact on precision than decreasing *N* has on reducing precision. However, it is likely that for very small values of *N*, the estimation error in branch lengths will grow rapidly.

An important exception to the overall trend was the age inferences for the crown eudicot node. The most common calibration strategy for this node has been to place a maximum bound or a highly informative prior on the age of this node, based on the absence of tricolpate pollen before the Barremian–Aptian boundary (~126 Ma) (Magallón and Castillo 2009; Sauquet et al. 2012; Massoni et al. 2015a; Foster et al. 2017). Additionally, many of the earliest-diverging eudicot lineages have relatively old fossils dating to the late Aptian (~113 Ma). These lines of evidence provide a narrow age bracket for the eudicot crown, often causing age estimates for the eudicot crown node to be necessarily highly precise. As a result, the limit in uncertainty of the fossil calibrations should be reached rapidly. Therefore, the age of the eudicot crown node is useful to evaluate in light of the finite-sites theory. We found that increasing the number of clock-subsets had essentially no effect on the uncertainty in the age estimate of this node. A very similar pattern was observed when using tightly constrained gamma calibration priors, and we expect that the general trend extends to other cases in which calibrated nodes have strongly constrained ages, for example when lognormal or exponential priors are chosen (Smith et al. 2010; Magallón et al. 2015).

Our results are especially important for analyses of genome-scale data sets. The size of phylogenomic data sets generally precludes molecular dating with computationally intensive phylogenetic software, such as BEAST (Bouckaert et al. 2014) or MrBayes (Ronquist et al. 2012), unless work-around methods are employed (Ho 2014). For example, some researchers have chosen to analyse each gene or data subset separately and then take the average of the results (Zeng et al. 2017). However, this methodology effectively assigns to each gene its own model of nucleotide substitution and its own clock model. Not only does this run the risk of severe overparameterization, but it also raises the question of how the estimates should be combined in a way that takes full account of estimation error. Another method is to apply data filtering to select only a subset of a data set, such as those that are the most clocklike (Jarvis et al. 2014) or the most informative (Tong et al. 2016).

In cases where data-filtering approaches are not feasible, less computationally intensive methods can be employed, such as the approximate-likelihood method of MCMCTREE. There are also non-Bayesian alternatives to phylogenomic dating, such as penalized likelihood (Sanderson 2002), that have been used to analyse large data sets (Zanne et al. 2014). Additionally, a number of rapid dating methods that can account for among-lineage rate heterogeneity without an explicit statistical model of branch-rate variation have been developed specifically for phylogenomic data sets (Kumar and Hedges 2016). Although these methods appear to have accuracy comparable to that of Bayesian methods, they cannot produce reliable estimates of the uncertainty in the inferred ages (Kumar and Hedges 2016). It is also unclear how well the results of these analyses will conform to the finite-sites theory.

## Conclusions

In this study, we have demonstrated that the finite-sites theory for molecular dating applies to a typical genome-scale data set from angiosperms, with the exception of nodes that have strong age constraints. In contrast with previous suggestions, the choice of strategy for assigning genes to clocks does not appear to be important. These results imply that the data set can be arbitrarily partitioned into a large number of clock-subsets, up to the point at which there is little marginal benefit in increasing the degree of clock-partitioning. However, we caution that all molecular date estimates should be critically interpreted to determine whether their precision is meaningful or not. To this end, the best approach is to identify the patterns of among-lineage rate heterogeneity in a data set and to apply a clock-partitioning scheme that appropriately captures this variation.

## Acknowledgements

The authors acknowledge the facilities and the technical assistance of the Sydney Informatics Hub at the University of Sydney and, in particular, access to the high-performance computing facility Artemis. This work was supported by the Research Training Program and the Australian Research Council [grant DP110100383 to S.Y.W.H.].

